# Cell-type-specific epigenomic variations associated with *BRCA1* mutation in pre-cancer human breast tissues

**DOI:** 10.1101/2020.08.24.265199

**Authors:** Yuan-Pang Hsieh, Lynette B. Naler, Sai Ma, Chang Lu

## Abstract

*BRCA1* germline mutation carriers are predisposed to breast cancers. Epigenomic regulations have been known to strongly interact with genetic variations and potentially mediate biochemical cascades involved in tumorigenesis. Due to the cell-type specificity of epigenomic features, profiling of individual cell types is critical for understanding the molecular events in various cellular compartments within complex breast tissue. Here we report cell-type-specific profiling of genome-wide histone modifications including H3K27ac and H3K4me3 in basal, luminal progenitor, mature luminal, and stromal cells extracted from pre-cancer *BRCA1* mutation carriers and non-carriers, conducted using a low-input technology that we developed. We discover that basal and stromal cells present the most extensive epigenomic differences between mutation carriers (*BRCA1*^*mut*/+^) and non-carriers (*BRCA1*^+/+^) while luminal progenitor and mature luminal cells are relatively unchanged with the mutation. Furthermore, the epigenomic changes in basal cells due to *BRCA1* mutation appear to facilitate their transformation into luminal progenitor cells. Our findings shed light on the pre-cancer epigenomic dynamics due to *BRCA1* mutation and how they may contribute to eventual development of predominantly basal-like breast cancer.

## Introduction

Mutations on tumor suppressor gene *BRCA1* have been strongly linked to increased risks to breast, ovarian and other cancers (Kuchenbaecker et al. 2017). However, how these genetic alterations trigger the molecular cascades that ultimately lead to the pathology of tumorigenesis remains unclear. Breast tissue contains both epithelial and stromal compartments and the former can be further divided into basal (BCs), luminal progenitors (LPs), and mature luminal (MLs) cells based on their surface markers that are indicative of their developmental lineage and/or location in the two epithelial layers of the mammary duct (Fu et al. 2014). These various cell types present characteristic gene expression patterns and epigenomic landscapes (Choudhury et al. 2013; dos Santos et al. 2015; Gascard et al. 2015; Pellacani et al. 2016). Breast tumors involving *BRCA1* germline mutation are predominantly basal-like (Foulkes 2004; Foulkes et al. 2004; Arnes et al. 2005). Recent results have suggested that *BRCA1*-associated basal-like breast cancers originate from luminal progenitor cells instead of basal stem cells (Lim et al. 2009a; Molyneux et al. 2010). Thus it is critical to understand how various cell types within breast tissue are affected by *BRCA1* mutation and how such dynamics in the cellular identity potentially contribute to tumorigenesis.

Epigenomic landscape plays a significant role in defining the cell state and mediating genetic factors into molecular cascades that are eventually involved in disease development. DNA sequence variation is known to impact epigenetic landscape, chromatin structures and molecular phenotypes via influencing the cis-regulatory elements such as promoters and enhancers (Kasowski et al. 2013; Kilpinen et al. 2013; McVicker et al. 2013). The changes in the epigenetic landscape may in turn alter gene expression and cellular phenotypes to promote cancer development. *BRCA1* mutation has been recently discovered to significantly alter epigenomic functional elements such as enhancers in our study using breast tissue homogenates (Zhang et al. 2019). However, due to predominant basal-like characteristic of *BRCA1*-associated tumors, cell-type-specific profiling of tissue samples is needed to decipher how each cell type within breast tissue is affected by the mutation and contributes to tumorigenesis.

In this study, we profile two important histone marks H3K4me3 and H3K27ac in a cell-type-specific manner in all four major cell types from pre-cancerous human breast tissue samples using a low-input ChIP-seq technology that we developed (MOWChIP-seq (Cao et al. 2015; Zhu et al. 2019)). We compare the data on *BRCA1* mutation carriers (*BRCA1*^*mut*/+^) and non-carriers (*BRCA1*^+/+^) and extract epigenomic features that separate the two groups. Such comparison reveals that the extent of epigenomic changes varies among the four cell types. These epigenomic alterations potentially change the cell state and lay the groundwork for future tumorigenesis.

## Results

Breast tissues from *BRCA1* mutation carriers (MUTs, n = 3) and non-carriers (NCs, n = 4) were collected during breast reduction or mastectomy surgery, dissociated, and sorted into the basal, luminal progenitor, mature luminal, and stromal cell (SC) types (Fig. 1a)(Chiang et al. 2012). We profiled H3K4me3 and H3K27ac using MOWChIP-seq with at least two technical replicates for each cell sample (Supplementary Tables S1 and S2). All samples had a fraction of reads in peaks (FrIP), normalized-strand correlation (NSC), and relative-strand correlation (RSC) that fell within ENCODE guidelines (Landt et al. 2012). H3K4me3 is an activating mark that is associated with transcriptional start sites of genes (Barski et al. 2007; Lauberth et al. 2013) and H3K27ac labels active enhancers (Creyghton et al. 2010). Our ChIP-seq datasets are highly correlated between technical replicates with an average Pearson correlation coefficient r of 0.962 for H3K4me3 and 0.950 for H3K27ac. We also observed very high genome-wide correlations among biological replicates in a group (MUTs or NCs), with an average r of 0.960 for H3K4me3 and 0.918 for H3K27ac (Fig. 1b). Generally, H3K4me3 is not a strong differentiating mark for separating MUTs and NCs. The correlation r between NCs and MUTs H3K4me3 data is high for all cell types (0.962 for BCs, 0.960 for LPs, 0.962 for MLs and 0.960 for SCs) (Supplementary Fig. S1). In contrast, when genome-wide H3K27ac is examined, many more differential peaks are observed between MUTs and NCs and among various cell types (Fig. 1c). BCs and SCs show large difference between MUTs and NCs (with an average r of 0.739 and 0.877, respectively). In comparison, LPs and MLs have similar H3K27ac profiles between MUT and NC (with an average r of 0.914 and 0.888, respectively).

**Figure 1.**
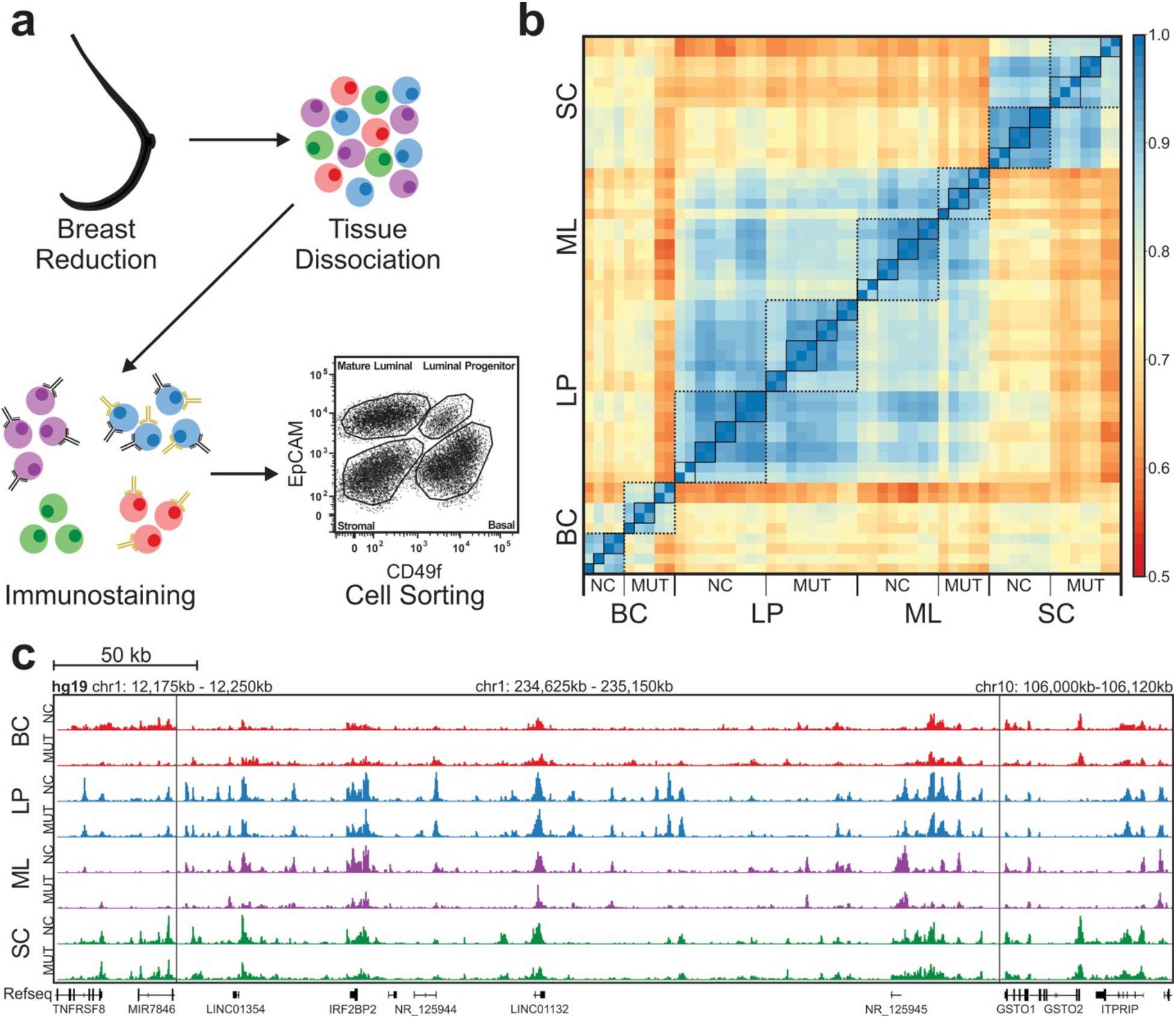
Cell-type-specific ChIP-seq data on human breast samples from BRCA1 mutation carriers (MUTs) and non-carriers (NCs). (a) Breast tissue samples were separated into the basal cells (BCs), luminal progenitor cells (LPs), mature luminal cells (MLs), and stromal cells (SCs) by FACS. (b) Pearson correlations among H3K27ac ChIP-seq data sets of various cell types from NCs and MUTs around promoter regions (TSS +/− 2kb). Each solid-line frame circles the technical replicates on one cell sample and each broken-line frame circles the data on a specific cell type. (c) Representative tracks of normalized H3K27ac signal for each of the cell types from NCs and MUTs. Three regions in the genome are presented and separated by vertical lines.

In terms of differences among various cell types, BCs, LPs, and MLs have very similar H3K4me3 profiles (average r of 0.926 among MUTs and 0.946 among NCs) and SCs show some differences from LPs and MLs (average r of 0.881 and 0.915 between SCs and LPs, 0.875 and 0.919 between SCs and MLs in MUT and NC, respectively). With H3K27ac data, LPs and MLs correlate with each other fairly well (average r = 0.832 and 0.872 in MUTs and NCs, respectively) while the other pairs have much lower correlation (with average r in the range of 0.668-0.794).

We carefully examined differentially modified H3K27ac peak regions (fold-change ≥ 2, FDR < 0.05) between MUTs and NCs (Fig. 2a). We found very few differential regions in LPs and MLs (518 and 2, respectively). However, there were a substantial number of differential peaks present in BCs (3,545) and a large number of different peaks in SCs (19,946). BCs had a mix of regions that showed either higher or lower H3K27ac signal in MUTs than in NCs (1,497 and 2,048, respectively), while the vast majority of differential regions in SCs (19,367 out of 19,946) had lower H3K27ac signal in MUT samples. We then compared the normalized H3K27ac signal at all peak regions (Fig. 2b). The median values were similar between NC and MUT patients in all epithelial cell types (with MUT values within ± 5% of NC ones), while there was a marked decrease in H3K27ac median signal in MUT SCs (by 13.5% compared to NC SCs).

**Figure 2.**
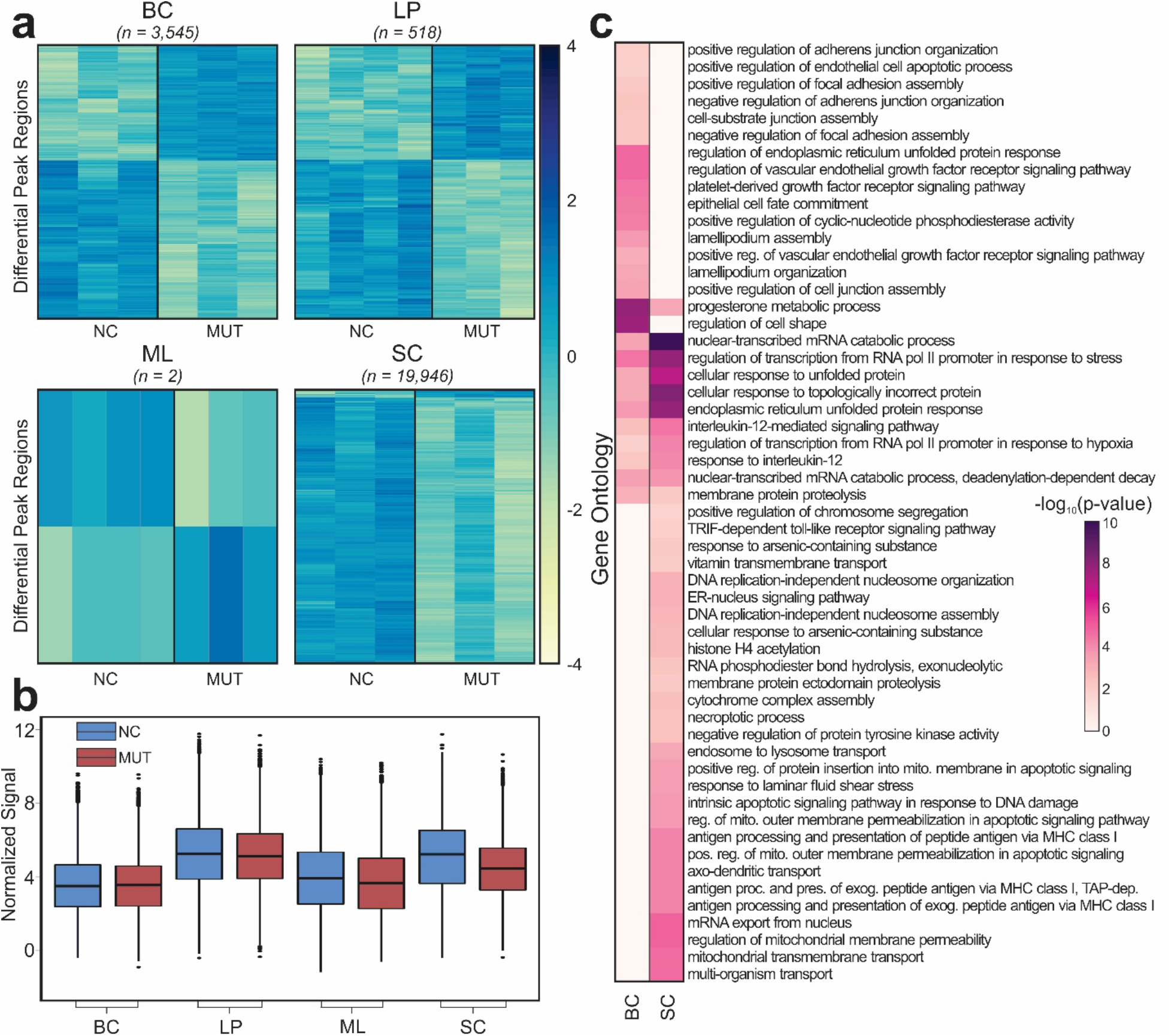
Differential H3K27ac peak regions between NCs and MUTs. (a) Heatmaps of differentially modified H3K27ac peak regions found to be significant (fold-change ≥ 2, FDR < 0.05) between NCs and MUTs. (b) Boxplots of normalized H3K27ac signal at all peak regions. (c) Gene ontology enrichment analysis using GREAT of significant differentially modified peak regions between NCs and MUTs.

The differentially modified H3K27ac regions were then mapped to their nearest genes (Supplementary Tables S3-S5). These differential regions were associated with 783, 3,972, and 12,160 genes in LPs, BCs, and SCs, respectively. We also conducted gene ontology enrichment analysis on these three cell types (Supplementary Table S6). The analysis on LPs did not bring out any GO terms. Thus our focus was on BCs and SCs which had the largest numbers of differential H3K27ac regions between NCs and MUTs (Fig. 2c). A number of BRCA1-associated processes, including progesterone metabolism (Katiyar et al. 2006; Ma et al. 2006; Calvo and Beato 2011; Davaadelger et al. 2019), RNA polymerase II transcription (Scully et al. 1997; Krum et al. 2003; Nair et al. 2016), and unfolded protein response (UPR) (Yeung et al. 2008), were enriched in both BCs and SCs. BRCA1 has been shown to inhibit progesterone signaling (Ma et al. 2006) and reduction in BRCA1 level has been shown to increase expression of GRP78, a key UPR regulating gene (Yeung et al. 2008). Moreover, BRCA1 is part of the RNA polymerase II holoenzyme (Scully et al. 1997). Ontologies related to apoptosis (Thangaraju et al. 2000; Andrews et al. 2002), antigen processing (Strickland et al. 2016; Green et al. 2017; Lu et al. 2019), and DNA damage response (Yoshida and Miki 2004; Wu et al. 2010) are only significant in SCs. BRCA1 has extensive association with apoptosis, including those due to endoplasmic reticulum stress that is related to UPR (Hedgepeth et al. 2015). For example, BRCA1 binding at the endoplasmic reticulum leads to a release of calcium that causes apoptosis. Furthermore, reduction in BRCA1 level has been shown to increase activation of CD8^+^ tumor-infiltrating lymphocytes. There are also several ontologies associated with DNA damage response significant in SCs. For instance, DNA replication-independent nucleosome assembly and organization can only occur with histone variant 3.3, which is part of the DNA repair pathway (Ahmad and Henikoff 2002; Frey et al. 2014). In addition, histone H4 acetylation also opens up the chromatin for easier access to damaged regions (Dhar et al. 2017). In contrast, we largely see ontologies associated with cell motility and adhesion in BCs. *BRCA1* mutations have been shown to increase cell motility in cancer cells (Yasmeen et al. 2008; Coene et al. 2011; Gau et al. 2015). However, epithelial cell fate commitment is also present only in BCs. Cells within the BC compartment have been previously shown to have the potential to differentiate into LPs (Holliday et al. 2018).

Next, we predicted enhancers present in each of the cell types for both NC and MUT samples (Fig. 3a). Enhancers were determined by finding H3K27ac^high^ regions that did not intersect with areas nearby transcription start sites (+/− 500bp from TSS). Using the NCs as the reference, MUTs covers 67%, 90%, 79%, and 29% of the NC enhancers in BCs, LPs, MLs, and SCs, respectively. The enhancers were then mapped to genomic regions (Fig. 3b, Supplementary Table S7). The most exaggerated differences due to *BRCA1* mutation were seen in BCs, including a 6.9% increase in the distal intergenic fraction and 12.3% decrease in the promoter vicinity fraction (i.e. < 2 kb from promoters). It is clear that *BRCA1* mutation plays a different role in enhancer activity that is unique to each cell type.

**Figure 3.**
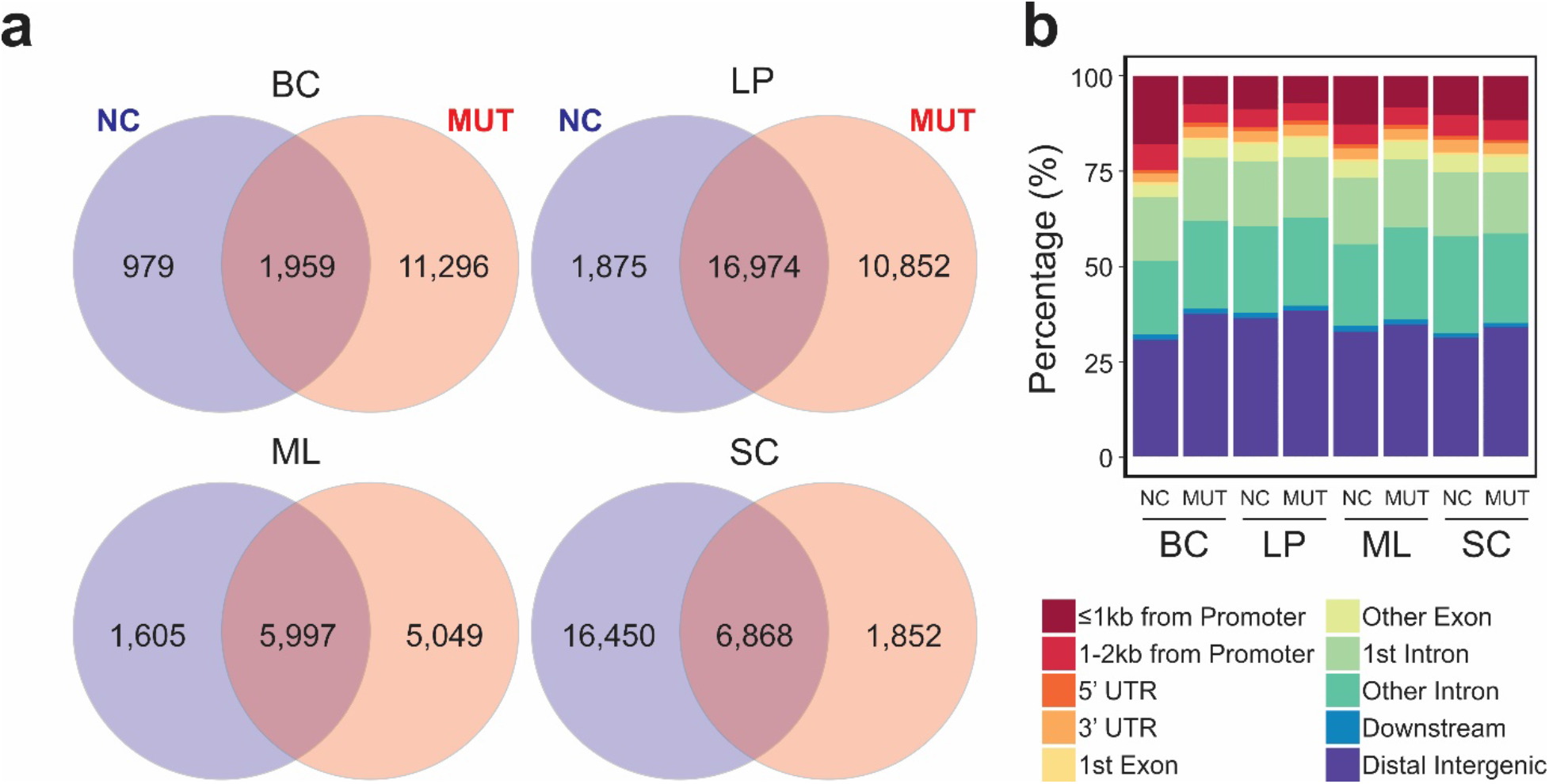
Enhancers predicted in various cell types from NCs and MUTs. (a) Venn diagrams of overlap between NC and MUT enhancers. (b) Genomic locations of the enhancers in various cell types.

Enhancer regions were then scanned to determine the transcription factor (TF) binding motifs significantly enriched in each cell population (Fig. 4, Supplementary Table S8). BCs had substantially more differentially enriched TF motifs than any other cell type, likely due to the smaller overlap between NC and MUT enhancers for BCs. In MUT samples, BCs primarily have many additional TFs, while SCs mainly lacked TFs. In all cases, over 50% of differentially enriched TFs were found to have no known link to *BRCA1* mutation. However, some TFs were found to be differentially enriched between MUT and NC in multiple cell types with known association with *BRCA1* mutation. These include *PAX5* (Branham et al. 2016) (BCs, MLs, and SCs), *CHOP* (Yeung et al. 2008) (BCs and SCs), *EGR1* (Lamber et al. 2010; Shin et al. 2013) (BCs and LPs), and *P73* (Ibrahim et al. 2010) (BCs and SCs). Some TFs found were not associated with *BRCA1* yet were associated with breast cancer. These include *EGR2* (Gregory et al. 2017; Li et al. 2017)(BCs, LPs, and SCs), *HOXC9* (Hur et al. 2014; Hur et al. 2016)(BCs, MLs, and SCs), *HSF1* (Gokmen-Polar and Badve 2016; Carpenter et al. 2017; Fujimoto et al. 2017)(BCs, LPs, and SCs), *NPAS2* (Lesicka et al. 2018)(BCs, LPs, and MLs), and *USF1* (Ramos et al. 2018)(BCs, LPs, and MLs). We then used PANTHER to classify the differential TFs in each cell type into pathways, and found that Gonadotropin-releasing hormone (GnRH) receptor pathway, Wnt Signaling pathway, apoptosis pathway, and p53 pathways were present in all four cell types. *BRCA1* has a key role in the Wnt signaling pathway(Li et al. 2010; Wu et al. 2012), regulates apoptotic responses(Thangaraju et al. 2000; Andrews et al. 2002), and has been shown to interact with *P53* (De Luca et al. 2013; Dong et al. 2015; Zhang et al. 2015; Peng et al. 2016). As for the GnRH pathway, while there is not a direct link to BRCA1, GnRH agonists have been shown to be effective in the treatment of breast cancer (Huerta-Reyes et al. 2019).

**Figure 4.**
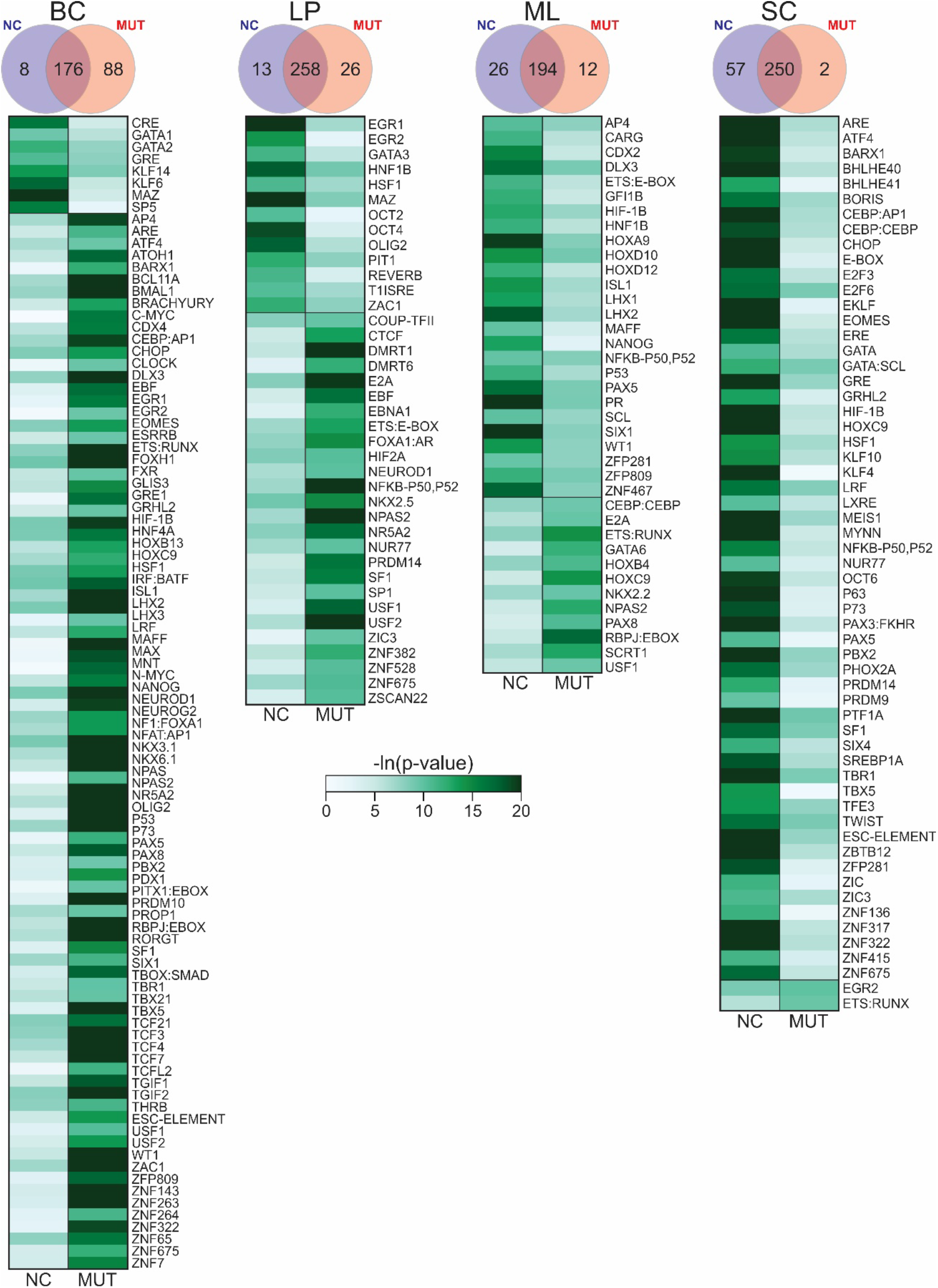
Heatmaps of motifs found to be significantly enriched in either NC or MUT but not both for various cell types. Color represents the level of significance. Venn diagrams of overlap between motifs enriched in NCs or MUTs are also presented.

Overall, we see the most significant epigenomic changes in BCs and SCs due to *BRCA1* mutation among the four cell types from these pre-cancer breast tissue samples, while much fewer variations were see in LPs. This is unexpected as LPs have been implicated as the driver in the onset of *BRCA1*-mutation associated breast cancer(Lim et al. 2009b). Thus we examine the possibility of basal cells differentiating into luminal cells, as proposed in previous literature (Holliday et al. 2018). First, we examined the cell-type specific genes identified for the three epithelial cell types (BCs, LPs, and MLs) in the literature (Pellacani et al. 2016). The expression of these genes largely defines the identity of specific cell types. By comparing NC and MUT BCs, we found that 6% (44/712) of the BC-specific genes experiences significant changes in H3K27ac state due to the mutation, compared to 1% (4/305) for LPs and 2% (23/444) for MLs (Supplementary Table S9). Of these differentially marked basal genes, 95% had lower H3K27ac signal in MUT, suggesting that there is primarily a loss of basal gene expression in MUT basal cells. In the same fashion, we also examined cell-type-specific TFs in the three cell types and how they vary due to the mutation. Enriched TFs in each cell population were predicted based on motif analysis of enhancers profiled using H3K27ac data (Ma et al. 2018). By examining NC samples, we extract 8, 55, and 6 cell-type-specific TFs (p-value < 0.0001) for BCs, LPs, and MLs, respectively (Fig. 5, Supplementary Table S10). In comparison, MUT BCs, LPs, and MLs preserved 6, 44, and 5 of these TFs, respectively. Interestingly, MUT BCs also enriched 28 of the LP-specific TFs and 3 of the ML-specific TFs, compared to MUT LP enriching 1 BC-specific TFs and 5 ML-specific TFs; and MUT ML enriching 2 BC-specific TFs and 5 LP-specific TFs. These results indicate that the BC state experiences more substantial change than LPs and MLs due to *BRCA1* mutation, consistent with the notion of BC differentiation into LPs.

**Fig. 5.**
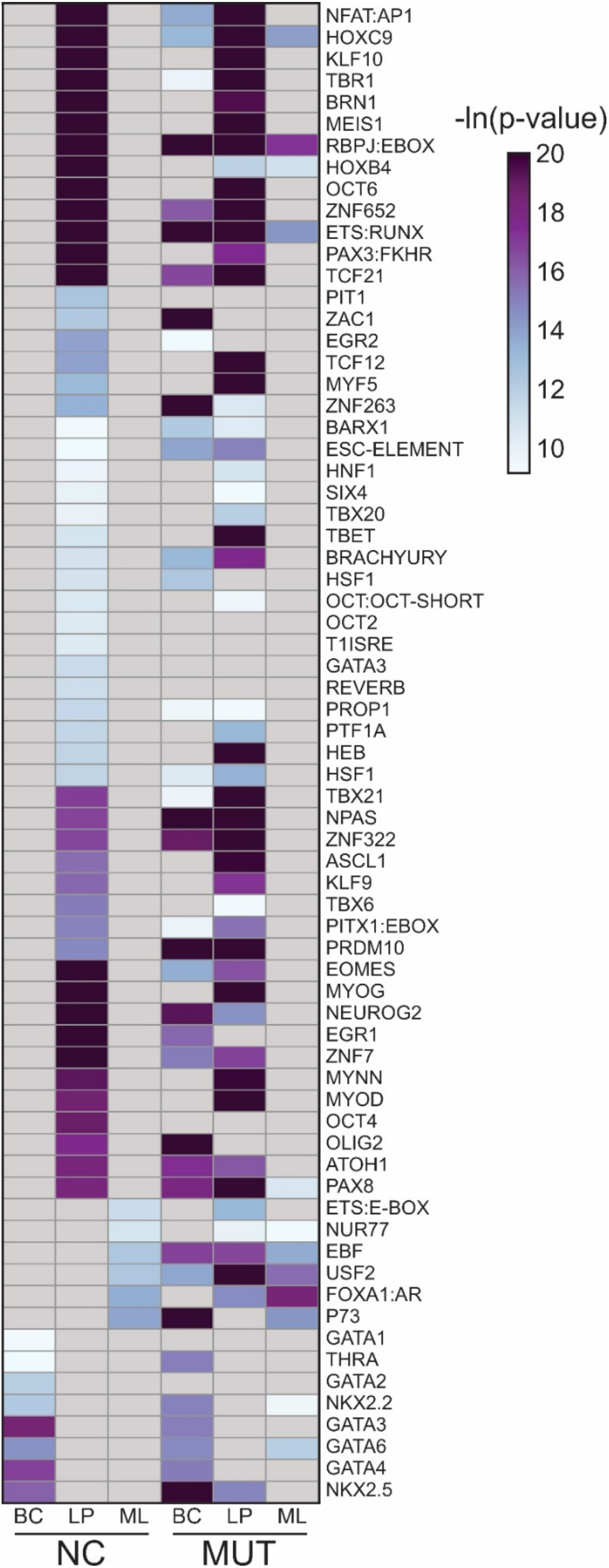
Heatmap of p-values for motifs found to be significantly enriched (p-value < 0.0001) in only one of three cell types (BCs, LPs, or MLs) from NCs and their enrichment in MUTs. Color represents the level of significance. All p-values that are insignificant are colored gray.

To further validate the possibility of the basal compartment being a significant contributor in *BRCA1*-mutation associated breast cancer, we determined the number of enhancers that were in proximity to 123 SNPs involved in ER-negative breast cancer as discovered by GWAS (Buniello et al. 2019) (Fig. 6). ER-negative breast cancer is a term that effectively covers both basal-like and triple-negative breast cancers (Rakha et al. 2008; Foulkes et al. 2010). In a cohort of 3,797 BRCA1 mutation carriers diagnosed with breast cancer, 78% had ER-negative breast cancers (Mavaddat et al. 2012). Overall, we saw increases in the percent of enhancers that were proximal to ER-negative SNPs in each cell type due to BRCA1 mutation, suggesting an overall increase in breast cancer risk. However, we see the largest increase (~55%) between NC and MUT BCs followed by SCs (~21%). This further supports that the BRCA1 mutation leads to profound epigenetic changes in BCs and that these changes have the potential to increase breast cancer risk.

**Fig. 6.**
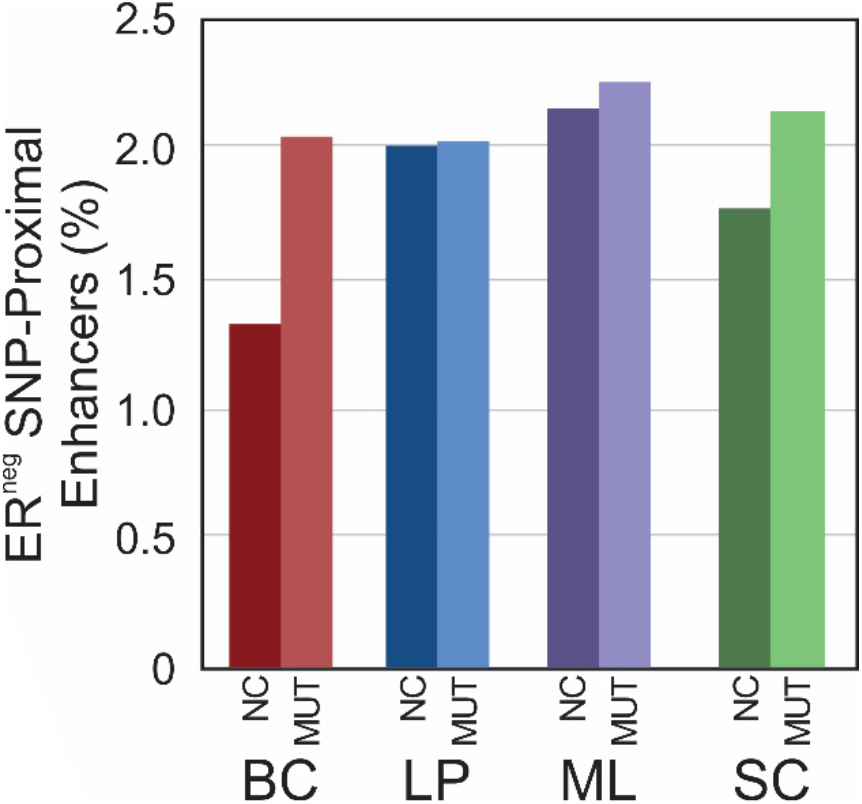
Percent of enhancers that are within +/− 150kb of SNPs significantly associated with ER-negative breast cancers through GWAS for all 4 cell types.

## Discussion

The interactions between genomics and epigenomics are well-recognized events. Gene mutation may alter epigenomic landscape in a significant way and such alternation may carry important implications on cancer development. Several lines of evidence support the feasibility of sorting out epigenomic differences between BRCA1 mutation carriers and non-carriers using a cell-type specific approach. First, we found very high correlations among biological replicates in both H3K4me3 and H3K27ac within either MUT or NC group. When we compare across the two groups (MUT vs. NC), H3K27ac is more differentiating than H3K4me3, which is consistent with previous findings by us (Ma et al. 2018) and others (Roadmap Epigenomics et al. 2015). Second, we compared our NC data with published results obtained by examining normal breast tissues pooled from multiple individuals (Pellacani et al. 2016). Each of the top 5 significantly enriched transcription factors in BCs, LPs, and MLs were also significantly enriched in our respective NC cell populations. Basal-associated transcription factors (Pellacani et al. 2016) such as TP53, TP63, STAT3, and SOX9 were also enriched in NC BCs. Similarly, luminal-associated transcription factors (Raouf et al. 2008; Chakrabarti et al. 2012; Theodorou et al. 2013) CEBPB, GATA3, ELF5, and FOXA1 were all significantly enriched in our NC LP and MLs. Third, we found that three members of the GATA family (GATA1, GATA2, and GATA3) were enriched in either NC BCs or LPs, but not in MUT epithelial cells (Fig. 4 and 5). This agrees with our earlier study conducted using breast tissue homogenates (Zhang et al. 2019). GATA3 is known to be critically involved in the regulation of luminal cell differentiation (Kouros-Mehr et al. 2006; Asselin-Labat et al. 2007). Finally, we also compared our cell-type-specific data with our published data on breast tissue homogenates obtained using a separate patient cohort and conventional ChIP-seq technology (Zhang et al. 2019) (Supplementary Fig. S2). The H3K27ac data taken using the breast tissue homogenate (mix) do not differentiate MUT and NC. The homogenate data show a similar degree of correlations with all individual cell types (Pearson correlation in the range of 0.741-0.890). This also underscores the importance for cell-type-specific profiling to pinpoint specific roles for each cell type. These comparisons suggest that although epigenomic differences exist among individual humans (McDaniell et al. 2010; Garg et al. 2018), careful cell-type-specific ChIP-seq profiling captures important genome-wide epigenomic differences due to *BRCA1* mutation.

Epigenetic profiles define cell identity by regulating cell-type-defining genes and transcription factors. The difference in the epigenomic landscape between *BRCA1* mutants and non-carriers may be important for explaining the high propensity of *BRCA1* mutation carriers for breast cancer. Our data on a sensitive mark H3K27ac are the most different in BCs and SCs when MUTs and NCs are compared. In comparison, very few changes were observed in LPs and MLs due to the mutation. Furthermore, our analysis of the cell-type-specific genes and TFs also reveal that MUT BCs resemble LPs. Such resemblance was in accordance with previous reports on the presence of LP-fated cells and bi-potent mammary stem cells in the basal compartment (Pal et al. 2017; Holliday et al. 2018). Thus we propose that the precancerous process within *BRCA1* mutation carriers may start with substantial epigenomic changes in basal cells among all epithelial cell types and these basal cells share similarity with luminal progenitor cells. These findings provide new insights into epigenomic factors involved in BRCA1 cancer biology.

## Methods

### Breast tissues

Breast tissues were obtained from adult female cancer-free *BRCA1* mutation carriers (MUT) or non-carriers (NC) who underwent cosmetic reduction of mammoplasty, diagnostic biopsies, or mastectomy. All procedures were approved by the University of Texas Health Science Center at San Antonio. The consent forms were signed by donors to approve the use of the tissue for breast cancer research. Genetic testing of *BRCA1* mutation was conducted by the hospital(Toland et al. 2018). Using previously published protocols(Zhang et al. 2017), fresh unfixed breast tissue was processed to generate single-cell suspension and the single cells were sorted into four fractions using FACS: EpCAM^−^CD49f^−^ stromal cells (SCs), EpCAM^low^CD49f^high^ basal cells (BCs), EpCAM^high^ CD49f^+^ luminal progenitor cells (LPs), and EpCAM^high^ CD49f^−^ mature luminal cells (MLs).

### Chromatin Shearing

The sonication process to generate chromatin fragments is similar to what we described in previous publications(Cao et al. 2015; Zhu et al. 2019). A sorted cell sample of a specific type (containing 100K to 3 million cells, depending on the cell type and sample) was centrifuged at 1600 g for 5 min at room temperature and washed twice with 1 ml PBS (4°C). Cells were resuspended in 1 ml of 1% freshly prepared formaldehyde in PBS and incubated at room temperature on a shaker for 5 min. Crosslinking was quenched by adding 0.05 ml of 2.5 M glycine and shaking for 5 min at room temperature. The crosslinked cells were centrifuged at 1600 g for 5 min and washed twice with 1 ml PBS (4°C). The pelleted cells were resuspended in 130 μl of the sonication buffer (Covaris, 10 mM Tris-HCl, pH 8.0, 1 mM EDTA, 0.1% SDS and 1x protease inhibitor cocktail (PIC)) and sonicated with 105 W peak incident power, 5% duty factor, and 200 cycles per burst for 16 min using a Covaris S220 sonicator (Covaris). The sonicated chromatin samples were shipped to Virginia Tech for MOWChIP-seq assay. The sonicated sample was centrifuged at 16,100 g for 10 min at 4 °C. The sheared chromatin in the supernatant was transferred to a pre-autoclaved 1.5 ml microcentrifuge tube (VWR). A fraction of the sonicated chromatin sample was mixed with IP buffer (20 Mm Tris-HCl, pH 8.0, 140 mM NaCl, 1 mM EDTA, 0.5 mM EGTA, 0.1 % (w/v) sodium deoxycholate, 0.1 % SDS, 1 % (v/v) Triton X-100, with 1% freshly added PMSF and PIC) to generate a MOWChIP sample containing chromatin from 50,000 cells with a total volume of 50 μl.

### MOWChIP-seq

We conducted MOWChIP-seq of the sonicated chromatin samples with 50,000 cells per assay for H3K27ac profiling and 10,000 cells per assay for H3K4me3 profiling, using protocols and microfluidic devices described in our previous publications(Cao et al. 2015; Zhu et al. 2019). We used anti-H3K27ac antibody (abcam, cat: ab4729, lot: GR323132-1) and anti-H3K4me3 antibody (Millipore, cat: 07-473, lot: 2930138) in these experiments.

### Data Quality Control

ChIP-seq data sets that had fewer than 10,000 called peaks were discarded. After quality control, the technical replicates of the same cell sample were combined for the data analysis. As the result, we obtained 3 biological replicates for MUT H3K27ac samples, 4 biological replicates for NC H3K27ac samples, 2 biological replicates for MUT H3K4me3 samples, and one biological replicate for the NC H3K4me3 sample. The fraction of reads in peaks (FrIP) was calculated using the number of mapped reads within peak regions divided by total mapped reads. Normalized-strand correlation (NSC) and relative-strand correlation (RSC) was calculated using phantompeakqualtools(Kharchenko et al. 2008; Landt et al. 2012).

### Data Processing

Unless otherwise mentioned, all data analysis was performed with Bash scripts or with R (The R Foundation) scripts in RStudio. Sequencing reads were trimmed using default settings by Trim Galore! (Babraham Institute). Trimmed reads were aligned to the hg19 genome with Bowtie(Langmead et al. 2009). Peaks were called using MACS2 (q < 0.05)(Zhang et al. 2008). Blacklisted regions in hg19 as defined by ENCODE were removed to improve data quality(Amemiya et al. 2019). Mapped reads from ChIP and input samples were extended by 100 bp on either side (250bp total) and a normalized signal was calculated.

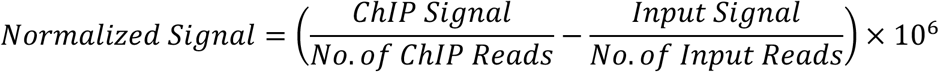

For Pearson’s correlation, the signal was calculated around the promoter region (TSS +/− 2kb) and plotted with the corr and levelplot functions. For visualization in IGV (Broad Institute), the signal was calculated in 100bp windows over the entire genome and output as a bigWig file.

### Differential Analysis

To determine peak regions with differential signal, the Bioconductor package DiffBind was used(Stark and Brown; Ross-Innes et al. 2012). A ‘majority-rules’ consensus peak set was generated for each experimental group and combined to make a master set for analysis. Peaks were considered to be valid if they were present in the majority of biological replicates. Counts were generated using default conditions and compared using the DESeq2 option. Normalized signal counts were extracted and plotted in heatmaps and boxplots using ggplot2(Wickham 2016). Gene ontology analysis was performed using the web-based tool GREAT 4.0.4(McLean et al. 2010) with default settings for hg19. For the SC analysis, the top 6,000 regions (by smallest FDR value) were used.

### Enhancers Analysis

To call enhancers, we considered H3K27ac^high^ regions that did not intersect with promoter regions to be enhancer regions. First, consensus H3K27ac peak sets were generated for NC and MUT samples for each cell type after determining the set of peak regions present in NC and/or MUT samples. Peak widths were expanded to be 1000 bp long (summit +/−500bp). Promoters were defined as TSS +/− 500 bp. Any H3K27ac 1kb regions that intersected with a promoter region was removed and the remaining regions were designated as enhancers. Motif analysis was performed to determine enriched transcription factor binding motifs among the enhancer regions with HOMER(Heinz et al. 2010) (with options ‒size 1000 ‒mask ‒p 16 ‒nomotif). Functional classification of transcription factors was performed using Panther v15.0.0(Mi et al. 2019). Enhancers were mapped to genomic regions with ChIPSeeker(Yu et al. 2015). Enhancers were considered associated with ER-negative SNPs (obtained from NHGRI-EBI GWAS Catalog(Buniello et al. 2019)) if the SNP was within 150kb up- or downstream.

## Data Availability

The ChIP-seq data sets are deposited in the Gene Expression Omnibus (GEO) repository with the following accession number: GSE148995. https://www.ncbi.nlm.nih.gov/geo/query/acc.cgi?acc=GSE148995 The following secure token has been created to allow review of record GSE148995 while it remains in private status: stypymsqdrmxzyp

## Acknowledgements

We thank Professors Rong Li and Xiaowen Zhang of George Washington University for providing the anonymous breast tissue samples and helpful discussion. This work was primarily supported by US National Institutes of Health (NIH) grant R33 CA214176 (C.L.) with additional support from NIH grants R01 CA243249 (C.L.), P30 CA012197 (C.L.), and a seed grant from Virginia Tech Institute for Critical Technology and Applied Science (C.L.).

## Author Contribution

C.L. designed and supervised the study. Y.-P.H. conducted the epigenomic profiling of the breast tissue samples. L.B.N. analyzed the data. S.M. helped with experimental work and data analysis. L.B.N. and C.L. wrote the manuscript. All authors proofread the manuscript and provided comments.

